# A sleep epidemic or enlightenment? A Bayesian approach to test the sleep epidemic hypothesis shows foragers have short and fragmented sleep compared to large scale societies

**DOI:** 10.1101/2020.09.16.299792

**Authors:** David R. Samson

**Affiliations:** University of Toronto, Mississauga

**Keywords:** mismatch, sleep, human evolution, circadian rhythm, mismatch, evolutionary medicine

## Abstract

Human sleep is linked with nearly every aspect of our health and wellbeing. The question whether and to what extent human sleep is in a state of evolutionary mismatch has gained recent attention from both clinical and biological science researchers. Here, I use a comparative Bayesian approach aimed at testing the sleep epidemic hypothesis – the idea that, due to labor demands and technological disruption, sleep-wake activity is negatively impacted in post-industrial, economically developed societies. In contrast to the expectations of the sleep epidemic hypothesis, when compared to both large and small-scale subsistence societies that rely on agriculture for subsistence, foragers were the shortest, least efficient sleeping group. Coupled with previous work demonstrating that foragers have stronger circadian rhythms compared to those sleeping in buffered environments, I present the sleep-rhythm trade-off hypothesis – that sleep duration, quality, and synchrony is driven by trade-offs between sleep security and comfort versus sleep site environmental exposure. One strategy to improve wellbeing of modern sleepers would be to focus on behavioral interventions that reduce desynchronizations of circadian rhythms, while holding the positive ground of safe, secure, and regulated sleep environments typical of economically developed societies.

## Introduction

In humans, sleep is paramount to nearly every facet of our health and well-being. Sleep determines our ability to think (cognition), feel (emotional regulation), socialize (interact positively with others), and preserve our health (immune strength, and cellular repair and maintenance) (Walker 2009). It follows that sleep is one of the most powerful predictors of performance and happiness over the life-course. Thus, understanding how a rapidly changing world, by way of globalization, migration, labor demands, technology and economic systems, is affecting our sleep – and how this past adaption mismatches with our changing world – is a critical challenge for the social and biological sciences in the 21^st^ century.

To this end, an interdisciplinary approach has emerged from anthropologists and sleep scientists to ‘take the sleep lab into the field’ to investigate how sleep in more “natural” environments (where artificial light and technology play a lesser role) and “artificial” urban environments (where artificial light and technology are more prevalent) differ in the ways in which they influence multiple dimensions of human health. This work has been timely, given the rapid rate of globalization and the exportation of cultural norms from developed economies influencing Indigenous populations across the globe.

Sleep deprivation in post-industrial, developed economies (i.e., WEIRD – Western, Educated, Industrialized, Rich, Democratic – nations (Henrich, Heine et al. 2010)) is argued to have been on the rise for the past five decades and to be reaching epidemic levels (Van Cauter and Knutson 2008, Roenneberg 2013). Growing technological disruption and innovation (Chang, Aeschbach et al. 2015), and high time-budget demands of the job market (Chatzitheochari and Arber 2009) have been suggested as primary drivers of the increase in sleep deprivation. This view, which I term here the ***sleep epidemic hypothesis,*** has led to increasing concerns for the general public (Colten and Altevogt 2006). Yet, the evidence in support of said public belief is scarce and contradictory (Lamote de Grignon Pérez, Gershuny et al. 2018). Whether sleep deprivation is in fact a growing problem has yet to be cross-culturally and comparatively explored and evaluated.

Furthermore, people in industrialized settings appear to be sleeping longer (Yetish, Kaplan et al. 2015, Samson, Crittenden et al. 2017, Samson, Crittenden et al. 2017). Indeed, compared to naturally sleeping groups within small-scale societies, individuals in developed economies appear to be experiencing longer duration and higher quality sleep via safer, more comfortable sleep sites. Yet, urban dwellers may be doing so at the cost of desynchronization (i.e., inconsistency and fragmentation) of circadian rhythms, which may lead to poor mental and physical outcomes. We call this alternative proposition, the ***sleep-rhythm trade-off hypothesis*** – that compared to naturally sleeping groups within small-scale societies, individuals in developed economies are experiencing long duration, higher quality sleep via safer, more comfortable sleep sites, at the cost of desynchronization (i.e., inconsistency and fragmentation) of circadian rhythms. Here, I test the sleep epidemic hypothesis and highlight alternative hypotheses, such as the sleep-rhythm trade-off hypothesis, for the aim of generating critical information for the development of interventions aimed at using evolutionary medicine to improving sleep, and thereby downstream health.

## Results

To analyse sleep globally I used Bayesian quadratic approximation. Two general approaches were adopted: (i) a parameter estimate of sleep duration of only actigraphy reported data (n = 28) and (ii) a data set that includes both actigraphy and PSG generated sleep duration (n = 59; Supplemental file 1). Based on the known reports of PSG and actigraphy sleep quotas I generated predictions of sleep duration and efficiency among primarily agricultural, hunter-gatherer (foragers), and non-forager small-scale subsistence societies (Table 1).

Assessing actigraphy generated sleep reports, the Bayesian quadratic approximation yielded Gaussian mean and standard deviation of 7.42 ± 0.18, with the credible interval of predicted values falling in-between 7.13 – 7.71 hours. The second, larger and methodologically mixed data set predicted a human sleep duration mean of 7.04 ± 0.11, with the credible interval as 6.86 – 7.23. Foragers were the shortest sleeping subsistence group (Table 1; Figure 1, panel (b)) with a predicted Gaussian mean and standard deviation of 6.56 ± 0.39, with the credible interval equalling 5.93 – 7.18 hour sleep duration. In addition, foragers were characterized by the least sleep efficiency generating a Gaussian mean and standard deviation of 77.87 ± 3.5, with a credible interval of predicted values between 72.3 – 83.5. Overall, foragers are predicted to sleep the least of any group compared to individuals that dwell within economically developing or developed societies that engage in agricultural practices.

**Table 1.** Predictions of night-time sleep duration and efficiency. Values were estimated using a Bayesian quadratic approximation approach. *Sigma* is equivalent to the predicted standard deviation for all categories combined. HDI is the Gaussian approximations for each parameter’s marginal distribution. The plausibility of each value of *mu* after averaging over the plausibility’s of each value of *sigma,* is given by the distributions mean and standard deviation. In this case, the percentile interval boundaries correspond to an 89% interval.

| **Subsistence strategy** | **Predicted mean and standard deviation** | **Highest Density Interval (HDI 89%)** |
| --- | --- | --- |
| Agriculture: sleep duration | 7.32 (0.30) | 6.84 – 7.81 |
| 4S-Forager: sleep duration | 6.56 (0.39) | 5.93 – 7.18 |
| 4S: sleep duration | 6.87 (0.23) | 6.50 – 7.24 |
| *Sigma*: sleep duration | 0.71 (0.13) | 0.51 – 0.91 |
| Agriculture: sleep efficiency | 86.92 (3.0) | 82.1 – 91.7 |
| 4S-Forager: sleep efficiency | 77.87 (3.5) | 72.3 – 83.5 |
| 4S: sleep efficiency | 80.07 (2.51) | 76.1 – 84.1 |
| *Sigma*: sleep efficiency | 7.48 (1.55) | 5.0 – 10.0 |

**Figure 1.**
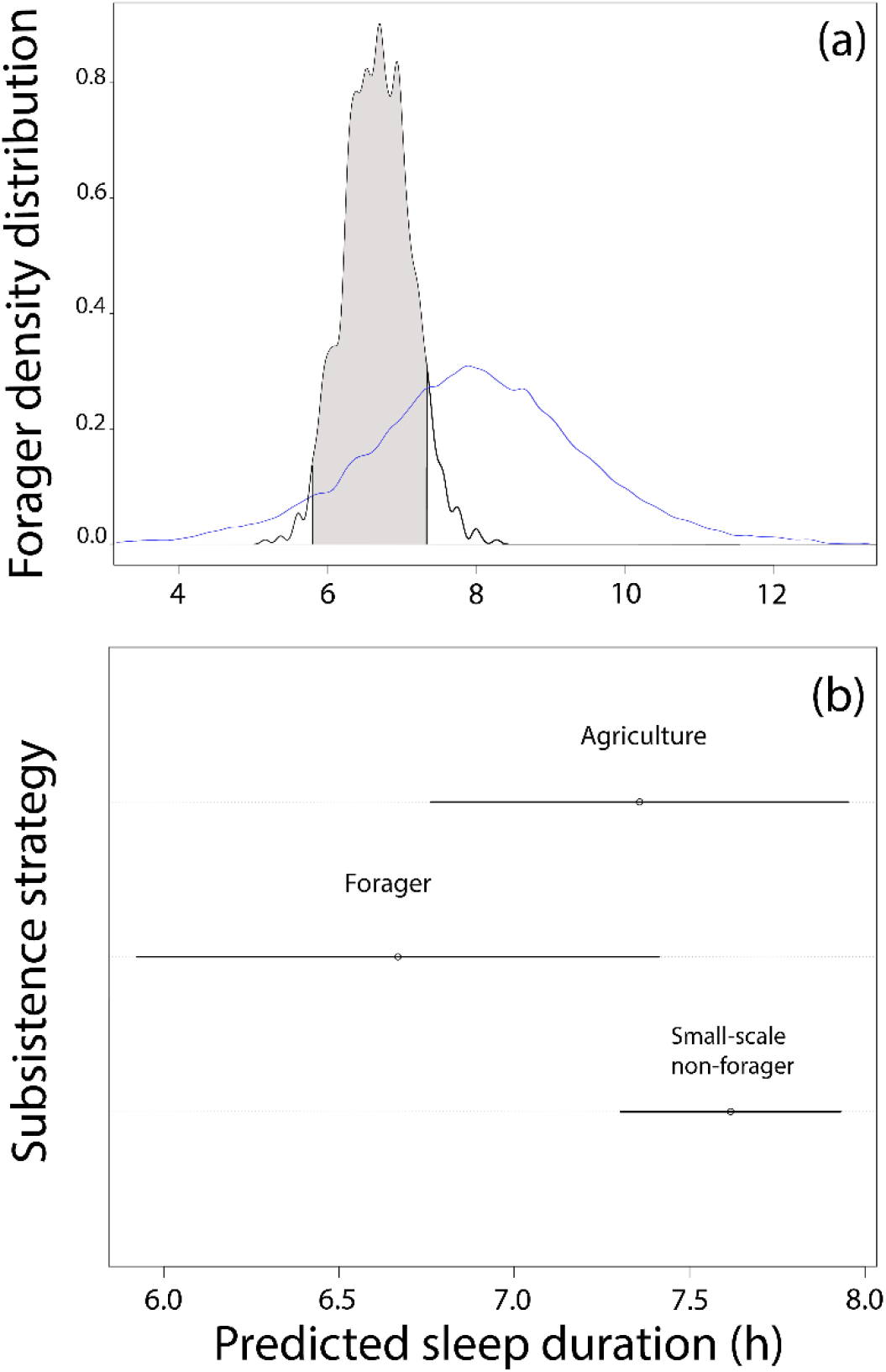
A Bayesian model using quadratic approximation to predict human sleep distributions. The parameters of interest are subsistence patterns influence of sleep duration. In panel (a) the blue line indicates the prior predictive distribution (10,000 simulations) that used a conservative μ of 8 (assuming the ideal 8-hour sleep period; see Materials and methods for more detail) and σ of 1 (assuming an idea range of between 7-9 hours sleep). Then, Bayesian updating considered every possible combination of values for sleep duration, taking into account each combination of parameters in the model to generate relative plausibility posterior probability distributions. In panel (a), for foragers this density distribution and the highest density interval (highlighted in grey) is shown to be left of the prior distribution. In panel (b) the model predicts the mean and 89% interval for each subsistence category, showing foragers to be characterized by the least duration of sleep, with none of the subsistence categories significantly extending beyond 8 hours of sleep.

## Discussion

By measures of sleep duration and efficiency, humans sleeping in modern, economically developed environments do not appear to be experiencing a sleep epidemic. The discovery that human forager sleep is especially short has critical implications for understanding modern human sleep and has important consequences for human health within a comparative, evolutionary lens.

Despite the well-documented disruptive effect of electricity and lighting on both developing (de la Iglesia, Fernández-Duque et al. 2015, Moreno, Vasconcelos et al. 2015, Pilz, Levandovski et al. 2018, Smit, Broesch et al. 2019) and developed societies (Chang, Aeschbach et al. 2015), with the advent of more secure, environmentally regulated sleep sites both sleep duration and efficiency have increased (Table 1). For optimized health, the American Association of Sleep Medicine recommends getting greater than 85% sleep efficiency (i.e., the time classified as asleep while in bed) (Ohayon, Wickwire et al. 2017). Societies that rely mostly on agriculture have the highest sleep efficiency (mean = 86.92), with non-forager 4S groups demonstrating less sleep efficiency (mean = 80.07%) and foragers showing the least (mean = 77.87%). Modern sleepers in more urban, economically developed environments may be sleeping deeper (i.e., greater proportion of slow-wave and REM) and more efficiently, but do so at the cost of desynchronization (i.e., inconsistency and fragmentation) of circadian rhythms, which may lead to poor downstream mental and physical health outcomes.

Consistent with those findings, Samson and colleagues (2017) recently reported that naturally sleeping (i.e., full-time exposure to daily variations in light and temperature) equatorial populations, such as small-scale agriculturalists in Madagascar and hunter-gatherers in Tanzania (Samson, Crittenden et al. 2017), have shorter, poorer sleep (i.e., more fragmented) than WEIRD populations, but stronger, more synchronized (i.e., greater amplitude and more consolidated) circadian rhythms. This suggests the intriguing possibility that there is an unexplored trade-off between circadian synchronization (i.e., a strong rhythm) and sleep duration and fragmentation. Importantly, short or fragmented (i.e., unconsolidated) sleep is associated with numerous negative health outcomes, including hypertension, obesity, chronic inflammation, kidney disease, and infertility (Miller and Cappuccio 2007, Bathgate, Edinger et al. 2015). On the other hand, circadian dysregulation (i.e., abnormal or de-amplified daily rhythms of sleep-wake activity, temperature, hormonal secretion) is associated with cancer, heart disease, and depressive disorders (Haus and Smolensky 2013, Grimaldi, Carter et al. 2016). Thus, there is evidence that although linked, sleep architecture (i.e., the distribution of non-rapid eye movement [NREM] and rapid eye movement [REM] sleep) and circadian rhythms have potentially independent effects. It stands to reason that transitioning to an environment that promotes sound sleep at the expense of circadian dysregulation is a profound trade-off, the implications of which have yet to be extensively investigated and highlights areas for future research.

Therefore, it appears that individuals in developed economies are experiencing long duration, higher quality sleep via safer, more comfortable sleep sites, at the cost of desynchronization (i.e., inconsistency and fragmentation) of circadian rhythms. The exploration of this idea – here called *the sleep-rhythm trade-off hypothesis* – is an urgent future research direction for sleep researchers to investigate.

Whether or not it is healthy for the average human to sleep 8 hours, the likelihood of this happening for the average human is low (Table 1, Figure 1). Although “paleo-sleep” may not be the healthiest sleep (Nunn and Samson 2019), given that hunter-gatherer subsistence strategies have been practiced since the emergence of genus *Homo* approximately 1.75 mya (Boehm 2012), it appears that there were evolutionary advantages to short sleep duration throughout human evolution (Samson and Nunn 2015). Critically, an evolutionary medicine perspective promotes the idea that natural selection operates on reproductive success, not optimized health (Stearns, Nesse et al. 2010). Therefore, with respect to human sleep duration and timing, clinical medicine can be correct if the focus is on health, whereas evolutionary anthropologists can be correct when describing the benefits to reproduction of specific sleep traits throughout evolutionary history. With growing understanding of the disruptive effects of light and temperature regulation of circadian regulation, one strategy to improve wellbeing of modern sleepers would be to focus on behavioral interventions that reduce desynchronizations of circadian rhythms, while holding the positive ground of safe, secure, and regulated sleep environments typical of economically developed societies.

## Materials and methods

For the comparative analysis of sleep quotas to apply Bayesian quadratic approximation, I generated a sleep database using both PSG and actigraphy reports from sleep studies performed in both the Cultural west and small-scale societies (Supplementary file 1). A major challenge for sleep anthropologists has been generating robust sleep measures for which to apply the comparative method to explore underlying patterns in sleep expression across societies. Since the discovery of sleep architecture in the 1950s (Dement and Kleitman 1953, Dement and Kleitman 1957), most sleep studies have been performed in the cultural West, in laboratory contexts, using the “gold standard” polysomnography (PSG) (Worthman 2008, Carskadon and Dement 2017). Given PSG uses electroencephalography (EEG) to measure brain states, it is highly accurate, yet it has known first night effects and may not capture the subjects normative sleep expression (Stone and Ancoli-Israel 2011) and is notoriously difficult to perform in dynamic field sites in ambulatory subjects (Samson, Yetish et al. 2016) and thus until the advent of reliable and cost effective wrist-worn accelerometer devices sleep remained measurable only in the laboratory (Worthman and Melby 2002). Actigraphy, otherwise known as accelerometry, has been a significant methodological advance for the study of the anthropology of sleep. It uses pieozoelectric film to measure activity by recording motion over a 24-hour cycles for extended periods (Cole, Kripke et al. 1992, Kripke, Hahn et al. 2010, Cellini, Buman et al. 2013, Samson, Yetish et al. 2016). Traditionally recorded in one-minute epochs the movement values are algorithmically scored to determine sleep wake states. Despite being vulnerable to overestimation of sleep states (de Souza, Benedito-Silva et al. 2003), actigraphy has been validated against PSG, demonstrating substantial improvements over subjective, self-reports of sleep (Lauderdale, Knutson et al. 2008).

### Statistical analysis

I used the R (Team 2016) rethinking package quadratic approximation ((McElreath 2012, McElreath 2018) to make inferences about the shape of the posterior distribution for sleep parameters. Subsistence strategy was classified as either predominantly reliant on agriculture, small-scale societies with differing mixtures of agriculture, horticulture, and pastoralism, or strict hunter-gatherer/forager communities. Sample size, which did not prove to be linked with sleep duration, was the number of participants used for each study. The model used is as follows:

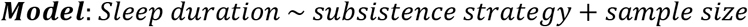

With respect to justification for the prior, Yetish and McGregor (2019) note that historical assumptions of desired sleep durations have informed the now ubiquitous conception of normative sleep being 8 hours – when in fact this may be historically linked to labor negotiations for sleep specific times as workers coped with the societal changes related to the industrial revolution and night-time production (Reiss 2017). The primary text for sleep clinicians, *Principles and Practices of Sleep Medicine,* states in the overview of normal human sleep that: “Most young adults report sleeping about 7.5 hours a night on weekday nights and slightly longer, 8.5 hours, on weekend nights,” (Carskadon & Dement 2017). Moreover, the National Sleep Foundation determined by panel vote how much sleep is required for overall health and well-being, suggesting that adult individuals (between 26-64 years old) should sleep 7-9 hours (Hirshkowitz et al 2015). Interestingly, averaging the ranges of the assessments in the latter two examples reifies the 8-hour assumption of the ideal human sleep period. In summary, these lines of evidence were used together to establish the prior of 8-hour

## ACKNOWLEDGMENTS

I am indebted to colleagues who reviewed, helped focus, and edit early versions of the manuscript: Gandhi Yetish, Eric Shattuck, Nancy Lai, Nickolas Tyler, and Charles Nunn. I am particularly grateful to my graduate students, Erica Kilius, Leela McKinnon, and Kaleigh Reyes for providing intellectual and structural feedback throughout the writing process.

## Notes

### Competing Interest Statement

The authors have declared no competing interest.

