## Supplemental file 1 for "A sleep epidemic or enlightenment? A Bayesian approach to test the sleep epidemic hypothesis shows foragers have short and fragmented sleep compared to large scale societies"

**Supplementary file 1. PSG and actigraphy based sleep studies across the globe.** Sleep duration and efficiency reported among primarily agricultural, hunter-gatherer (foragers), and non-forager small-scale subsistence societies.

| Country | Population | Method | Primary subsistence strategy | Sleep duration (h) | Sleep efficiency | Sample size | Reference |
|---|---|---|---|---|---|---|---|
| USA | Maryland | PSG | Agriculture | 6.49 | | 30 | (Feinberg et al 1967) |
| USA | New York | PSG | Agriculture | 6.38 | | 10 | (Kahn et al 1970) |
| USA | Florida | PSG | Agriculture | 6.28 | 91 | 10 | (Williams et al 1972) |
| Scotland | Edinburgh | PSG | Agriculture | 7.58 | | 14 | (Březinová 1975, Březinová 1976) |
| Switzerland | Geneva | PSG | Agriculture | 8.17 | 95.5 | 40 | (Gaillard 1978) |
| USA | Gainsville | PSG | Agriculture | 6.70 | 87 | 109 | (Berry & Webb 1985) |
| USA | San francisco | PSG | Agriculture | 6.88 | 91.0 | 23 | (Naifeh et al 1987) |
| USA | Pittsburgh | PSG | Agriculture | 6.08 | 80.0 | 19 | (Hoch et al 1988) |
| USA | New York | PSG | Agriculture | 6.17 | 86.9 | 40 | (Schiavi et al 1988) |
| USA | California | PSG | Agriculture | 6.13 | | 24 | (Bonnet 1989) |
| Netherland | Groningen | PSG | Agriculture | 6.00 | | 28 | (Dijk et al 1989) |
| USA | Pittsburgh | PSG | Agriculture | 6.88 | 87.8 | 24 | (Brendel et al 1990) |
| USA | Pittsburgh | PSG | Agriculture | 5.45 | 73.7 | 105 | (Brendel et al 1990) |
| USA | Washington | PSG | Agriculture | 6.42 | | 24 | (Vitiello et al 1990) |
| Germany | Mannheim | PSG | Agriculture | 6.54 | 89.7 | 51 | (Lauer et al 1991) |
| USA | Pittsburgh | PSG | Agriculture | 6.63 | 85.1 | 64 | (Monk et al 1991) |
| USA | Houston | PSG | Agriculture | 5.81 | 86.9 | 186 | (Hirshkowitz et al 1992) |
| USA | Pittsburgh | PSG | Agriculture | 6.62 | 85.1 | 66 | (Monk et al 1992) |
| USA | Pittsburgh | PSG | Agriculture | 6.19 | 82.6 | 50 | (Hoch et al 1994) |
| USA | Chicago | PSG | Agriculture | 7.13 | 72.2 | 16 | (Nofzinger et al 1995) |

| | | | | | | | |
|---|---|---|---|---|---|---|---|
| Switzerland | Zurich | PSG | Agriculture | 7.16 | 89.6 | 16 | (Landolt et al 1996) |
| USA | Washington | PSG | Agriculture | 6.40 | 84 | 113 | (Vitiello et al 1996) |
| USA | Pittsburgh | PSG | Agriculture | 7.23 | 93.9 | 61 | (Ehlers & Kupfer 1997) |
| Israel | Haifa | PSG | Agriculture | 5.70 | 80.6 | 25 | (Haimov & Lavie 1997) |
| Italy | Parma | PSG | Agriculture | 7.46 | 89.5 | 40 | (Parrino et al 1998) |
| USA | California | PSG | Agriculture | 6.85 | 90.5 | 73 | (Rao et al 1999) |
| USA | Dallas | PSG | Agriculture | 6.62 | | 23 | (Fulton et al 2000) |
| USA | Pittsburgh | PSG | Agriculture | 6.94 | 95.2 | 100 | (Carrier et al 2001) |
| Canada | Montréal | PSG | Agriculture | 7.85 | 93.8 | 30 | (Gaudreau et al 2001) |
| France | Lyon | PSG | Agriculture | 7.35 | 93.3 | 36 | (Nicolas et al 2001) |
| Australia | Melbourne | PSG | Agriculture | 7.56 | 94.9 | 14 | (Crowley et al 2002) |
| Italy | Bologna | Actigraphy | Agriculture | 8.10 | 94.0 | 282 | (Natale et al 2009) |
| USA | Chicago | Actigraphy | Agriculture | 7.02 | 90.0 | 126 | (Carnethon et al 2016) |
| USA | California | Actigraphy | Agriculture | 6.55 | 83.0 | 495 | (Yoon et al 2003) |
| Sri Lanka | Colombo | Actigraphy | Agriculture | 6.00 | 85.0 | 133 | (Schokman 2018) |
| Tanzania | Hadza | Actigraphy | 4S-Forager | 6.22 | 67.0 | 21 | (Samson et al 2017a) |
| Namibia | San | Actigraphy | 4S-Forager | 6.66 | 83.7 | 175 | (Yetish et al 2015) |
| Congo | BaYaka | Actigraphy | 4S-Forager | 6.06 | 67.0 | 33 | (Samson et al 2020a; Unpublished work) |
| Northern Namibia | Himba | Actigraphy | 4S | 5.34 | 65.0 | 27 | (Prall et al 2018) |
| Bolivia | Tsimané | Actigraphy | 4S | 6.63 | 83.7 | 62 | (Yetish et al 2015) |
| Madagascar | Malagasy | Actigraphy | 4S | 6.55 | 70.0 | 75 | (Samson et al 2017b) |
| Argentina | Toba/Qom | Actigraphy | 4S | 6.90 | | 120 | (de la Iglesia et al 2015) |

| | | | | | | | |
|---|---|---|---|---|---|---|---|
| Argentina | Toba/Qom | Actigraphy | 4S | 7.73 | | 21 | (de la Iglesia et al 2015) |
| Haiti | Fondwa | Actigraphy | 4S | 7.00 | 89.0 | 19 | (Knutson 2014) |
| Mozambique | Tengua | Actigraphy | 4S | 7.23 | 78.5 | 19 | (Beale et al 2017) |
| Mozambique | Milange | Actigraphy | 4S | 7.28 | 83.3 | 58 | (Beale et al 2017) |
| Guatemala | Mayan | Actigraphy | 4S | 6.60 | 80.0 | 34 | (Samson et al 2020b; Unpublished work) |
| Brazil | Chico Mendez Amazonian Reserve | Actigraphy | 4S | 8.10 | | 28 | (Moreno et al 2015) |
| Brazil | Chico Mendez Amazonian Reserve | Actigraphy | 4S | 8.35 | | 64 | (Moreno et al 2015) |
| Brazil | Bombas | Actigraphy | 4S | 8.96 | | 114 | (Pilz et al 2018) |
| Brazil | Aria Branca | Actigraphy | 4S | 9.17 | | 268 | (Pilz et al 2018) |
| Brazil | Sao Roque | Actigraphy | 4S | 7.92 | | 15 | (Pilz et al 2018) |
| Brazil | Corrego do Franco | Actigraphy | 4S | 8.77 | | 15 | (Pilz et al 2018) |
| Brazil | Mamas | Actigraphy | 4S | 7.90 | | 15 | (Pilz et al 2018) |
| Brazil | Morro do Fortunato | Actigraphy | 4S | 8.65 | | 15 | (Pilz et al 2018) |
| Brazil | Peixoto dos Botinhas | Actigraphy | 4S | 7.66 | | 15 | (Pilz et al 2018) |
| Vanuatua | Tanna Island | Actigraphy | 4S | 7.88 | 82.7 | 15 | (Smit et al 2019) |
| Vanuatua | Tanna Island | Actigraphy | 4S | 7.42 | 80.3 | 15 | (Smit et al 2019) |

*****Note: The first anthropological study of sleep in a 4S was performed in Papua New Guinea (Siegmund et al 1998) but is omitted here due to reported 24-hour total sleep durations and not night time sleep duration.

### SUPPLEMENTARY MATERIAL LITERATURE CITED


Beale AD, Pedrazzoli M, Bruno da Silva BG, Beijamini F, Duarte NE, et al. 2017. Comparison between an African town and a neighbouring village shows delayed, but not decreased, sleep during the early stages of urbanisation. *Scientific reports* 7: 5697

Berry D, Webb WB. 1985. Sleep and cognitive functions in normal older adults. *Journal of Gerontology* 40: 331-35

Bonnet M. 1989. Infrequent periodic sleep disruption: effects on sleep, performance and mood. *Physiology & behavior* 45: 1049-55

Brendel DH, Reynolds III C, Jennings J, Hoch C, Monk T, et al. 1990. Sleep stage physiology, mood, and vigilance responses to total sleep deprivation in healthy 80-year-olds and 20-year-olds. *Psychophysiology* 27: 677-85

Březinová V. 1975. The number and duration of the episodes of the various EEG stages of sleep in young and older people. *Electroencephalography and clinical neurophysiology* 39: 273-78

Březinová V. 1976. Duration of EEG sleep stages in different types of disturbed night sleep. The Fellowship of Postgraduate Medicine

Carnethon MR, De Chavez PJ, Zee PC, Kim K-YA, Liu K, et al. 2016. Disparities in sleep characteristics by race/ethnicity in a population-based sample: Chicago Area Sleep Study. *Sleep medicine* 18: 50-55

Carrier J, Land S, Buysse DJ, Kupfer DJ, Monk TH. 2001. The effects of age and gender on sleep EEG power spectral density in the middle years of life (ages 20–60 years old). *Psychophysiology* 38: 232-42

Carskadon MA, Dement WC. 2017. Normal human sleep: an overview  In *Principles and Practice in Sleep Medicine*, ed. MH Kryger, T Roth, WC Dement, pp. 15-24. Philadelphia, PA: Elsevier Saunders

Cellini N, Buman MP, McDevitt EA, Ricker AA, Mednick SC. 2013. Direct comparison of two actigraphy devices with polysomnographically recorded naps in healthy young adults. *Chronobiology international* 30: 691-98

Cole RJ, Kripke DF, Gruen W, Mullaney DJ, Gillin JC. 1992. Automatic sleep/wake identification from wrist activity. *Sleep* 15: 461-69

Crowley K, Trinder J, Kim Y, Carrington M, Colrain IM. 2002. The effects of normal aging on sleep spindle and K-complex production. *Clinical neurophysiology* 113: 1615-22

de la Iglesia H, Fernández-Duque E, Golombek DA, Lanza N, Duffy JF, et al. 2015. Access to electric light is associated with shorter sleep duration in a traditionally hunter-gatherer community. *Journal of biological rhythms* 30: 342-50

de Souza L, Benedito-Silva AA, Pires MLN, Poyares D, Tufik S, Calil HM. 2003. Further validation of actigraphy for sleep studies. *Sleep* 26: 81-85

Dement W, Kleitman N. 1953. Regularly occurring periods of eye motility, and concomitant phenomena, during sleep. *Science* 118: 273-74

Dement W, Kleitman N. 1957. Cyclic variations in EEG during sleep and their relation to eye movements, body motility, and dreaming. *Electroencephalogryaphic Clinical Neurophysiology* 9: 673-90

Dijk DJ, Beersma DG, Bloem GM. 1989. Sex differences in the sleep EEG of young adults: visual scoring and spectral analysis. *Sleep* 12: 500-07

Ehlers C, Kupfer D. 1997. Slow-wave sleep: do young adult men and women age differently? *Journal of sleep research* 6: 211-15

Feinberg I, Koresko RL, Heller N. 1967. EEG sleep patterns as a function of normal and pathological aging in man. *Journal of psychiatric research* 5: 107-44

Fulton MK, Armitage R, Rush AJ. 2000. Sleep electroencephalographic coherence abnormalities in individuals at high risk for depression: a pilot study. *Biological psychiatry* 47: 618-25

Gaillard J-M. 1978. Chronic primary insomnia: possible physiopathological involvement of slow wave sleep deficiency. *Sleep* 1: 133-47

Gaudreau H, Carrier J, Montplaisir J. 2001. Age-related modifications of NREM sleep EEG: from childhood to middle age. *Journal of sleep research* 10: 165-72

Haimov I, Lavie P. 1997. Circadian Characteristics of Sleep Propensity Function in Healthy Elderly: A Comparison With Young Adults. *Sleep* 20: 294-300

Hirshkowitz M, Moore CA, Rando K, Karacan I. 1992. Polysomnography of adults and elderly: sleep architecture, respiration, and leg movement. *Journal of clinical neurophysiology: official publication of the American Electroencephalographic Society* 9: 56-62

Hoch CC, Dew MA, Reynolds III CF, Monk TH, Buysse DJ, et al. 1994. A longitudinal study of laboratory- and diary-based sleep measures in healthy “old old” and “young old” volunteers. *Sleep* 17: 489-96

Hoch CC, Reynolds CF, Houck PR, Verran JA, Wolanin MO. 1988. Sleep patterns in Alzheimer, depressed, and healthy elderly. *Western journal of nursing research* 10: 239-56

Kahn E, Fisher C, Lieberman L. 1970. Sleep characteristics of the human aged female. *Comprehensive Psychiatry* 11: 274-78

Knutson KL. 2014. Sleep duration, quality, and timing and their associations with age in a community without electricity in haiti. *American Journal of Human Biology* 26: 80-86

Kripke DF, Hahn EK, Grizas AP, Wadiak KH, Loving RT, et al. 2010. Wrist actigraphic scoring for sleep laboratory patients: algorithm development. *Journal of sleep research* 19: 612-19

Landolt H-P, Dijk D-J, Achermann P, Borbely AA. 1996. Effect of age on the sleep EEG: slow-wave activity and spindle frequency activity in young and middle-aged men. *Brain research* 738: 205-12

Lauderdale DS, Knutson KL, Yan LL, Liu K, Rathouz PJ. 2008. Self-reported and measured sleep duration: how similar are they? *Epidemiology (Cambridge, Mass.)* 19: 838-45

Lauer CJ, Riemann D, Wiegand M, Berger M. 1991. From early to late adulthood changes in EEG sleep of depressed patients and healthy volunteers. *Biological psychiatry* 29: 979-93

Monk TH, Reynolds III CF, Buysse DJ, Hoch CC, Jarrett DB, et al. 1991. Circadian characteristics of healthy 80-year-olds and their relationship to objectively recorded sleep. *Journal of Gerontology* 46: M171-M75

Monk TH, Reynolds III CF, Machen MA, Kupfer DJ. 1992. Daily social rhythms in the elderly and their relation to objectively recorded sleep. *Sleep* 15: 322-29

Moreno CRdC, Vasconcelos S, Marqueze EC, Lowden A, Middleton B, et al. 2015. Sleep patterns in Amazon rubber tappers with and without electric light at home. *Scientific reports* 5: 14074

Naifeh KH, Severinghaus JW, Kamiya J. 1987. Effect of aging on sleep-related changes in respiratory variables. *Sleep* 10: 160-71

Natale V, Plazzi G, Martoni M. 2009. Actigraphy in the assessment of insomnia: a quantitative approach. *Sleep* 32: 767-71

Nicolas A, Petit D, Rompre S, Montplaisir J. 2001. Sleep spindle characteristics in healthy subjects of different age groups. *Clinical Neurophysiology* 112: 521-27

Nofzinger EA, Reynolds CF, Thase ME, Frank E, Jennings JR, et al. 1995. REM sleep enhancement by bupropion in depressed men. *The American journal of psychiatry*

Parrino L, Boselli M, Spaggiari MC, Smerieri A, Terzano MG. 1998. Cyclic alternating pattern (CAP) in normal sleep: polysomnographic parameters in different age groups. *Electroencephalography and Clinical Neurophysiology* 107: 439-50

Pilz LK, Levandovski R, Oliveira MA, Hidalgo MP, Roenneberg T. 2018. Sleep and light exposure across different levels of urbanisation in Brazilian communities. *Scientific reports* 8: 11389

Prall SP, Yetish G, Scelza BA, Siegel JM. 2018. The influence of age-and sex-specific labor demands on sleep in Namibian agropastoralists. *Sleep health* 4: 500-08

Rao U, Poland RE, Lutchmansingh P, Ott GE, McCracken JT, Lin K-M. 1999. Relationship between ethnicity and sleep patterns in normal controls: implications for psychopathology and treatment. *Journal of Psychiatric Research* 33: 419-26

Samson DR, Crittenden AN, Mabulla AI, Mabulla AZP, Nunn CL. 2017a. Hadza sleep biology: evidence for flexible sleep-wake patterns in hunter-gatherers. *American Journal of Physical Anthropology* 162: 573-82

Samson DR, Kilius E, Lew-Levy S, Sarma M, Gettler LT, Boyette A. 2020a. Shifting sleep ecologies among foragers: An intra-community comparison of village and forest sleep of BaYaka foragers from the Congo.

Samson DR, Manus MB, Krystal AD, Fakir E, Yu JJ, Nunn CL. 2017b. Segmented sleep in a nonelectric, small-scale agricultural society in Madagascar. *American Journal of Human Biology*: 1-13

Samson DR, McKinnon L, Nunn CL, Nepomnaschy PA. 2020b. Male sleep is shorter and more fragmented than female sleep in a semi-electric, non-industrial, rural population of Kaqchikel Maya.

Samson DR, Yetish G, Crittenden AN, Mabulla I, Mabulla AZP, Nunn CL. 2016. What is segmented sleep? Actigraphy field validation for daytime sleep and nighttime wake. In *Journal of Sleep Research*, pp. 16. Journal of Sleep Research

Schiavi RC, Schreiner-Engel P, White D, Mandeli J. 1988. Pituitary-gonadal function during sleep in men with hypoactive sexual desire and in normal controls. *Psychosomatic Medicine*

Schokman AS. 2018. *Measures of Sleep Duration and Quality in Sri Lanka*. University of Sydney

Siegmund R, Tittel M, Schiefenhovel W. 1998. Activity monitoring of the inhabitants in Tauwema, a traditional melanesian village: Rest/activity behaviour of Trobriand islanders (Papua New Guinea). *Biological Rhythm Research* 29: 49-59

Smit AN, Broesch T, Siegel JM, Mistlberger RE. 2019. Sleep timing and duration in indigenous villages with and without electric lighting on Tanna Island, Vanuatu. *Scientific reports* 9: 1-16

Stone KL, Ancoli-Israel A. 2011. Actigraphy  In *Principles and practice of sleep medicine*, ed. MH Kryger, T Roth, C William, pp. 1668-75. St. Louis, Missouri: Elsevier Saunders

Vitiello MV, Larsen LH, Moe KE, Borson S, Schwartz RS, Prinz PN. 1996. Objective sleep quality of healthy older men and women is differentially disrupted by nighttime periodic blood sampling via indwelling catheter. *Sleep* 19: 304-11

Vitiello MV, Prinz PN, Williams DE, Frommlet MS, Ries RK. 1990. Sleep disturbances in patients with mild-stage Alzheimer's disease. *Journal of gerontology* 45: M131-M38

Williams RL, Karacan I, Hursch CJ, Davis CE. 1972. Sleep patterns of pubertal males. *Pediatric research* 6: 643

Worthman CM, ed. 2008. *After dark: the evolutionary ecology of human sleep*. Oxford: Oxford University Press. 291-313 pp.

Worthman CM, Melby MK. 2002. Toward a Comparative Developmental Ecology of Human Sleep  In *Adolescent sleep patterns: biological, social, and psychological influences*, ed. MA Carskadon, pp. 69-117. Cambridge: Cambridge University Press

Yetish G, Kaplan H, Gurven M, Wood B, Pontzer H, et al. 2015. Natural Sleep and Its Seasonal Variations in Three Pre-indistrial Societies. *Current Biology* 25: 1-7

Yoon IY, Kripke DF, Youngstedt SD, Elliott JA. 2003. Actigraphy suggests age-related differences in napping and nocturnal sleep. *Journal of sleep research* 12: 87-93
